# Cutaneous alternating current stimulation can cause a phasic modulation of speech perception

**DOI:** 10.1101/2023.12.26.573341

**Authors:** Jules Erkens, Ram K. Pari, Marina Inyutina, Mathieu Marx, Florian H. Kasten, Benedikt Zoefel

## Abstract

Segregating important stimuli from distractors is crucial for successful speech perception. Neural activity synchronized to speech, also termed “neural speech tracking”, is thought to be instrumental for this purpose. However, the relative contribution of neural tracking of targets and distractors for speech perception in a setting with multiple competing speakers remained unclear. In 61 human participants, we used transcranial alternating current stimulation (tACS) to manipulate neural tracking of two simultaneously presented sequences of rhythmic speech while participants attended to one of them. A random temporal relationship between speech streams allowed us to disentangle effects of tACS on target and distractor processing, and to examine their combined effect on a behavioural measure of speech perception. We found that the phase relation between tACS and both target and distracting speech modulated word report accuracy to a similar degree. This effect was observed during bilateral tACS over auditory regions, the inferior frontal gyrus (IFG) and, importantly, in a control group that received near-identical cutaneous stimulation but ∼50% reduced brain stimulation. These results imply that, although neural tracking of both target- and distracting speech causally modulates speech perception, the cutaneous stimulation that goes along with tACS can cause phase effects in speech perception that resemble those for conventional tACS. Our finding illustrates the urgent need to control for cutaneous stimulation in tACS studies.

**Significance Statement:** Neural activity synchronised to speech (“neural speech tracking”) plays an important role in speech perception, yet its exact role in multi-speaker scenarios remains underexplored. We here use tACS to manipulate neural tracking in such scenarios. We find that neural tracking of both target and distracting speech causally modulates speech perception, irrespective of whether tACS is applied to target auditory cortex, inferior frontal gyrus, or in a control condition designed to produce similar cutaneous stimulation, but ∼50% reduced direct brain stimulation. Although our results demonstrate a causal role of neural tracking for speech perception in multi-speaker scenarios, they also illustrate the urgent need to control for effects of cutaneous stimulation in future tACS work on speech perception and beyond.

## Introduction

Neural entrainment is the process of synchronizing ongoing neural activity to rhythmic stimuli (Lakatos et al., 2008; Schroeder & Lakatos, 2009). Evidence has accumulated that neural entrainment structures human perception, particularly that of speech (Henry & Obleser, 2012; Horton et al., 2013; Kasten et al., 2024; Kösem et al., 2018; Peelle et al., 2013; Peelle & Davis, 2012; Ten Oever & Sack, 2015; Zoefel, 2018). A related term that avoids implicit and debated (Haegens & Zion Golumbic, 2018; Obleser & Kayser, 2019; Zoefel, ten Oever, et al., 2018) assumptions about underlying neural mechanisms (such as the involvement of neural oscillations) is “neural speech tracking”, a term we adopt in this article.

Brain stimulation methods have been a major asset to this field of research, allowing causal interference about the role of neural tracking in speech perception that would be impossible with brain imaging alone (Herrmann et al., 2016; Kasten & Herrmann, 2022; Vosskuhl et al., 2018; Zoefel & Davis, 2017). One such method is transcranial alternating current stimulation (tACS), a non-invasive stimulation method that is believed to modulate neural activity by subthreshold polarization of the resting membrane potential (Antal & Paulus, 2013; Bland & Sale, 2019). By varying the tACS waveform relative to the speech stimulus and testing for consequences in perception, tACS has been used successfully to modulate speech processing (Riecke et al., 2018; van Bree et al., 2021; Wilsch et al., 2018; Zoefel et al., 2018, 2020; for review, see Preisig, 2024).

In one of the few studies that applied tACS to modulate neural tracking in multi-speaker scenarios, Keshavarzi et al. (2021) independently varied the timing of tACS relative to target- and distracting speech, showing that the perception of target speech is disrupted when its envelope was conveyed through tACS at a phase shift other than 0° (i.e. if it differed from the acoustic envelope). The tACS waveform conveying the distractor envelope also modulated target perception, with a 180° shift (relative to target) leading to most accurate perception. This finding implies that neural tracking of both target- and distracting speech causally modulates speech perception. However, as the tracking of target speech and that of the distractor were modulated in separate conditions, their combined effects could not be investigated. In the current study, we used rhythmically spoken speech to simultaneously vary the phase of ongoing tACS relative to both target- and distracting speech. A random temporal relationship between targets and distractors allowed us to disentangle their effects in the same trials (Figure 1A): For a given phase relation between tACS and target speech, the distractor was presented at all possible tACS phases, averaging out any phasic effect of the latter (and vice versa). Thus, we could quantify causal effects of neural tracking of target- and distracting speech, both combined and separately (Figure 1B).

**Figure 1.**
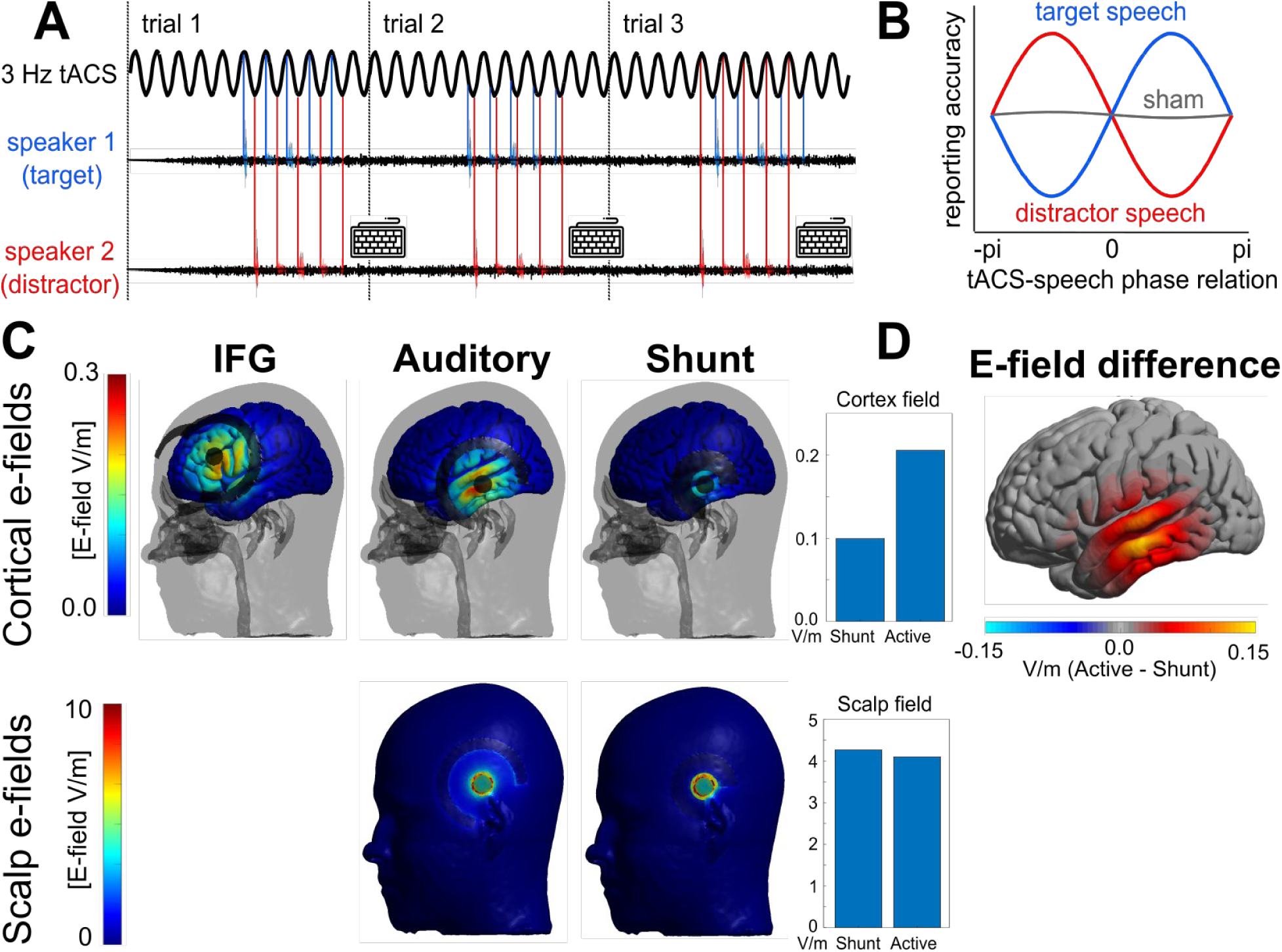
**A**: Experimental setup. Target- and distracting speech streams were presented rhythmically at specific phases of ongoing 3-Hz tACS. Both the phase relation between tACS and the two speech streams and the relative delay between the two were random and varied between trials. Consequently, the tACS phase for target speech was uncorrelated with that for distracting speech, and their effects could be disentangled. After each five-word sequence, participants were asked to type in the middle three words from the target speech (keyboard symbol). **B**: Hypothetical effect of tACS phase on task performance. We tested whether the accuracy of reporting target words depends on tACS phase relative to target speech (red) or distracting speech (blue), and whether these effects differ in magnitude. Whereas the phase relation between tACS and target speech that leads to enhanced target perception typically differs across participants (+pi/2 in the example shown), we expected that the same phase relation amplifies distracting information and therefore leads to poor target perception when applied to distracting speech. This hypothesis predicts opposite phase relation between the two conditions (target vs distracting speech) within participants. **C:** TACS montages and an estimate of the electric field they induce in the cortex (top) and scalp (bottom), based on a template brain and the average stimulation intensity used (2.1 mA peak to peak for Auditory and Shunt, 1.6 mA for IFG). Auditory and Shunt: Center electrode at positions T7 and T8 of the conventional EEG 10-10 system. IFG: Individualised tACS montages targeting pars opercularis of the IFG. The electric field in the scalp is more than one order of magnitude stronger than that in the brain, even for active tACS (note the difference in scale between the two rows). Scalp e-fields are only shown for auditory and shunt montages, as these are contrasted directly. The bars illustrate peak field strengths for these two montages (average of the 2000 voxels with the strongest e-fields; other N do not significantly alter the relative difference in peak field strength between montages). The Shunt montage minimises cortical stimulation despite near-identical cutaneous stimulation as compared to the Auditory montage. **D:** Difference in cortical electric field between Auditory and Shunt montages.

The above-mentioned studies used tACS to target activity in auditory cortical regions. However, successful speech perception involves a wide network of brain regions (Hickok & Poeppel, 2015), and the representation of target- and distracting speech becomes more distinct at higher levels of the auditory hierarchy (Zion Golumbic et al., 2013). We therefore additionally tested whether neural tracking in the inferior frontal gyrus (IFG) plays a causal role for speech perception in a multi-speaker setting.

During tACS, most of the applied current is shunted by the skin. It has been shown that transcutaneous stimulation of peripheral nerves can cause effects in the motor system that resemble those observed during cortical stimulation, suggesting that cutaneous stimulation, and subsequent relay to sensorimotor systems, might underlie some of the tACS effects (Asamoah et al., 2019b). However, tACS entrains single-neuron activity in non-human primates even with somatosensory input blocked (Vieira et al., 2020), and the extent to which cutaneous stimulation explains tACS effects is unclear. Cutaneous origins of tACS effects on speech perception remain untested. However, they seem to be supported by recent studies showing that vibrotactile pulses can modulate speech perception (Guilleminot et al., 2023; Guilleminot & Reichenbach, 2022), despite the fact that tactile stimulation can modulate neural speech tracking without concomitant changes in speech perception (Riecke et al., 2019). In three different experimental groups, we applied bilateral tACS to target auditory primary cortex or inferior frontal gyrus (Figure 1C), as well as a montage over auditory regions designed to produce equivalent cutaneous stimulation, but ∼50% reduced brain stimulation (Neri et al., 2020, Figure 1D). We tested whether and how tACS in these three groups modulates speech perception in the multi-speaker setting described above.

## Materials and Methods

### Participants

Sixty-one participants (Mean age 25.66, St. Dev. 3.12, 31 women) were recruited for the experiment. They provided informed consent under a process approved by the Comité de Protection des Personnes (CPP, protocols 2020-A02707-32 and 2021-A01446-35). All participants were native French speakers, reported having normal hearing, and had no history of neurological disease or any other exclusion criteria for tACS. They were divided into three different groups that only differed in their electrode montages and corresponding target region. Each group underwent a single stimulation condition (auditory, IFG, or shunt/control). Groups were tested in the order “auditory”, “IFG”, “control/shunt”, and included twenty-three, sixteen and twenty-two participants, respectively.

### Stimuli and Experimental Design

Stimuli consisted of sequences of one-syllable French words presented at a comfortable level using in-ear headphones (ER2 Earphones, Etymotic Research Inc., USA). Words were randomly selected from a pool of ∼500 words that were spoken individually to a metronome at ∼2 Hz (inaudible to participants). Each trial consisted of two sequences of five words, embedded in speech-shaped background noise and time-compressed to 3 Hz using the change tempo function of Audacity (version 3.2.0, Audacity Team, 2021). The two sequences were presented simultaneously, but with a variable lag relative to each other (Figure 1A), described in detail below. The signal-to-noise ratio between speech and background noise was 5 dB, determined in pre-tests to achieve ∼50 % task accuracy and kept identical across participants. One of the two-word sequences was spoken by a male speaker, and the other one by a female speaker. The speech-shaped noise was computed from the average spectrum of all words (male and female).

The first and last words were ‘pause’ in each sequence and irrelevant for participants’ task. We made this choice as the first and last words had reduced overlap with the other speech stream, and would thus be easier to perceive. Participants were instructed to attend to one of the two speech streams (male or female), and ignore the other. After each sequence, participants were asked to report the middle three words from the five-word sequence they had attended, using a standard computer keyboard. Participants performed ten blocks of this task; the first two blocks familiarized the participant with the task (performance was not included in the analysis), followed by eight experimental blocks. Each block consisted of 36 trials of approximately 9.3 seconds each. The identity of the target speaker alternated between male and female between blocks. Participants were encouraged to guess or fill in partial words if they were uncertain. After each block, participants were asked to rate their sensations from the stimulation at a scale that ranged from “no sensations at all” (equivalent to 0 points) to “strong sensations” (equivalent to 10 points). Five (out of sixty-one) participants did not fill in the sensation questionnaire. In between blocks, participants were allowed to take breaks at their leisure. The task was presented using Matlab (The MathWorks Inc., 2022. MATLAB version: 9.7 (R2019b), Natick, Massachusetts: The MathWorks Inc. https://www.mathworks.com) on a Windows 10 PC.

For both speech streams separately, the five words were presented so that the perceptual centre (Scott, 1998) of each word aligned at one out of six different phases of ongoing 3-Hz tACS (Figure 1A). These six phases covered one complete 3-Hz cycle, i.e., they varied from 0° to 300° in steps of 60° (with 360° being omitted as it is equivalent to 0°), or from 0 ms to 277.78 ms in steps of 55.56 ms, respectively. The tACS phase for each trial was selected pseudo-randomly, and separately for both speech streams. Consequently, the phase (and temporal) delay between target- and distracting speech was also random, ranging from complete overlap (in-phase) to no overlap (anti-phase). Our design created 36 different combinations of tACS-target and tACS-distractor phase relations. Within each block, each of these combinations was presented once. For the sham blocks, the same 36 target-distractor phase combinations were used to maintain the relative timing of the two speech streams. This resulted in a total of 6 (blocks) x 36 = 216 tACS trials (36 per phase and attended/distractor condition) and 2 (blocks) x 36 = 72 sham stimulation trials (36 per attended/distractor condition as tACS phase is not defined for sham).

### Electric stimulation

TACS was administered using two battery-driven stimulators (DC-Stimulator MR, Neuroconn GmbH, Ilmenau, Germany), one for each hemisphere. To assure synchronization between current and sound, stimulators and participants’ headphones were driven by the output of the same high-quality sound card (Fireface UCX, RME, Germany; sampling rate 44100 Hz). The tACS electrode configuration consisted of a bilateral ring setup that produced reliable modulation of speech perception and neural oscillatory activity in previous works (van Bree et al., 2021; Zoefel, Allard, et al., 2019). Electrodes were attached with adhesive, conductive ten20 paste (Weaver and Company, Aurora, CO, USA). The electrode montage was prepared in such a way that the skin impedances on both sides of the head were comparable and never exceeded 10 kOhm. During six experimental blocks, tACS was applied continuously for the entire block. At the start and end of the block, tACS was faded in and out using the first- and second half of an eleven second Hanning window, respectively. The two remaining blocks served as sham blocks, for which current was faded in and out using the same Hanning window (i.e. within eleven seconds), but without a 3-Hz component and without stimulation for the remainder of the block. This was done to simulate the typical sensations caused by tACS, which are most pronounced at the beginning of the stimulation. The order of stimulation and sham blocks was randomized, and the two sham blocks always consisted of one male and one female target speaker block. These stimulation parameters were used for all three electrode montages.

For tACS of auditory regions (Figure 1C, top), centre electrodes with a diameter of 20 mm were attached at positions T7 and T8 of the conventional electroencephalography (EEG) 10-10 system. The outer ring electrodes, with a band diameter of 13 mm (outer/inner diameter: 100/75 mm), were placed around these centre electrodes, with 25 mm distance on all sides. A quarter circle was cut out of the outer rings as to not overlap with the ear. For tACS of IFG (Figure 1C, bottom), we targeted pars opercularis. As stimulation of IFG is less established for the current design, tACS electrode locations were optimised for individual participants. Using structural T1 and T2 MRI scans, individual head models were created and then segmented into different tissues using the CHARM algorithm of SIMNIBS 4.0 (Puonti et al., 2020). A custom search tree algorithm was used to determine the montage that produces the highest maximum field strength (based on the 5% voxels with the strongest E-field in the target region) in the pars opercularis. We restricted possible tACS electrode locations to those in the 10-10 system and at midpoint positions in-between. The electrode shapes and distances between them were identical to those of auditory tACS. The montage for cutaneous stimulation (Figure 1D, top) was identical to that for auditory tACS, but the outer ring was replaced with one of 22 mm band diameter (outer/inner diameter: 75/30 mm) and 5 mm distance from the centre electrode. Based on electric field modelling, this electrode design was chosen to produce near-identical cutaneous stimulation but ∼50% reduced cortical stimulation (Figure 1D, bottom). As the close distance between electrodes increases the risk of gel bridges during cutaneous stimulation, we verified that impedances were not implausibly low (i.e. close to 0 kΩ). For all stimulation montages, current intensity was adjusted to a level comfortable to the participant, with an average intensity of 2.1 mA (peak to peak; std: 0.8 mA), 1.6 mA (std: 0.6 mA) and 2.1 mA (std: 0.4 mA) for auditory, IFG, and cutaneous stimulation, respectively.

### Analysis: Word report accuracy

To evaluate participants’ accuracy in reporting words from the target sequence, their written responses and the target words were first converted to a phonological representation using the phonemizer Python plugin (Bernard & Titeux, 2021). Responses and targets were then compared using Levenshtein distance, i.e. the minimum number of substitutions, deletions and insertions necessary to turn one word into the other (Levenshtein, 1966). The Levenshtein distance is typically normalized by dividing its absolute value by the length of target or response word (de la Higuera & Mico, 2008). In this way, however, the same absolute Levenshtein distance can produce multiple normalized outcomes, depending on the length of these words. To avoid this issue, we used an alternative approach that was validated in simulations during pre-tests. This approach “corrected” participants’ responses by regressing single-trial Levenshtein distances on the length of corresponding target words, and subtracting the explained variance. All subsequent analyses were then performed using the residuals from this regression analysis. As each trial consisted of three target words, participants entered up to three words after each trial, and these might not follow the original order. We therefore calculated Levenshtein distances for each combination of target and response words within a trial and selected the combination with the highest accuracy. Note (1) that the same procedure was applied to the sham condition which was later compared with tACS (see next section) and (2) that it does not bias accuracy specifically for some tACS phases and therefore cannot have produced spurious phase effects.

### Analysis: Comparison with sham

The random phase relationship between targets and distractors allowed us to disentangle their causal contributions to word report accuracy: For a given phase relation between tACS and target speech, the distractor was presented at all possible tACS phases, and any phasic effect the latter produces would cancel out on average (and vice versa). However, due to the nature of the design, target speech always preceded the distractor for some tACS phases, and always followed it for other tACS phases. These differences in *global* timing directly affected the first and last words of the sequence, as these were either preceded or followed by a word from the other speech stream or not, depending on the tACS phase. These words were therefore excluded from the quantification of word report accuracy. For the middle three words, indirect effects of global timing (“*contextual* masking”) were present but could be controlled for, as they were equally prevalent during tACS and sham stimulation (see Results). For each relative timing between speech streams (corresponding to the 36 target-distractor phase combinations described above) and for single participants in each of the three groups, we first calculated the trial-averaged accuracy (corrected Levenshtein distance as described in the previous section) in the sham condition. From the accuracy in each tACS trial, we then subtracted the average accuracy in the sham condition obtained for the respective target-distractor combination. This approach allowed us to isolate effects of tACS phase (which should only be present in the tACS condition) from generic effects of presentation timing. It also allowed us to express performance relative to a baseline (sham) condition.

### Analysis: Phasic modulation of word report accuracy

To quantify whether tACS had a phasic effect on word report accuracy (corrected for target word length and compared with sham) in any of three experimental groups, we applied a statistical approach that was shown to be particularly sensitive at detecting such effects (Zoefel et al., 2019). We used a regression model to test how strongly tACS phase (sine- and cosine-transformed) predicts single-trial measures of performance. This model yields two regression coefficients for individual participants (one for sine, one for cosine predictor) that together reflect the magnitude of phasic modulation. The two coefficients were extracted for tACS phase effects on target and distracting speech, respectively.

To test for reliable phase effects, we compared the coefficients from the individual regression models that included (sine and cosine-transformed) tACS phase as a predictor with those obtained from an intercept-only regression model, using an F-test (function “coefTest” in MATLAB; example code can be found in Zoefel et al., 2019). This comparison yielded one p-value per participant. This p-value is low if the regression model that includes tACS phase better predicts behavioural outcomes than a model that only includes an intercept term (but no tACS phase). Individual p-values were combined to a group level p-value using Fisher’s method. We performed this analysis separately for the two conditions (target and distractor tACS), as well as for a combined regression model which uses the tACS phase from both conditions (i.e. two sine and two cosine-transformed phases) as predictor for behavioural outcomes. We then compared the fit of these models using Fisher’s transformation of the correlation coefficient (Fisher, 1928).

For comparison, we constructed a distribution of regression coefficients that would be obtained under the null hypothesis of no phasic modulation of speech perception. This null distribution was obtained by randomly assigning tACS phases to single trials before re-computing regression coefficients, separately for each participant. This procedure was then repeated 1000 times, yielding 1000 average regression coefficients that would have been obtained in the absence of a tACS effect (which was abolished by the randomization of phases).

To contrast tACS effects between montages and conditions (target vs distractor modulation), we subjected regression coefficients to a mixed model ANOVA (between subjects: montages auditory/IFG/control, within subjects: target/distractor).

Participants typically differ in their “best” or “worst” tACS phases, i.e., those leading to most or least accurate perception, respectively (Riecke et al., 2015; van Bree et al., 2021; Zoefel, Davis, et al., 2019). We extracted participants’ best phases from the individual regression models. These phases correspond to the peak of a sine function fitted to word report accuracy as a function of tACS phase (Figure 2A). As this method relies on data from all tACS phases rather than defining best phase exclusively based on maximum performance (Erkens et al., 2020, 2021; Riecke et al., 2018), this method is less affected by task-unrelated noise in the data. We used Rayleigh’s test for non-uniformity to test whether participants’ best tACS phases were distributed non-uniformly (“circ_rtest” from Circular Statistics Toolbox for MATLAB; Berens, 2009).

**Figure 2.**
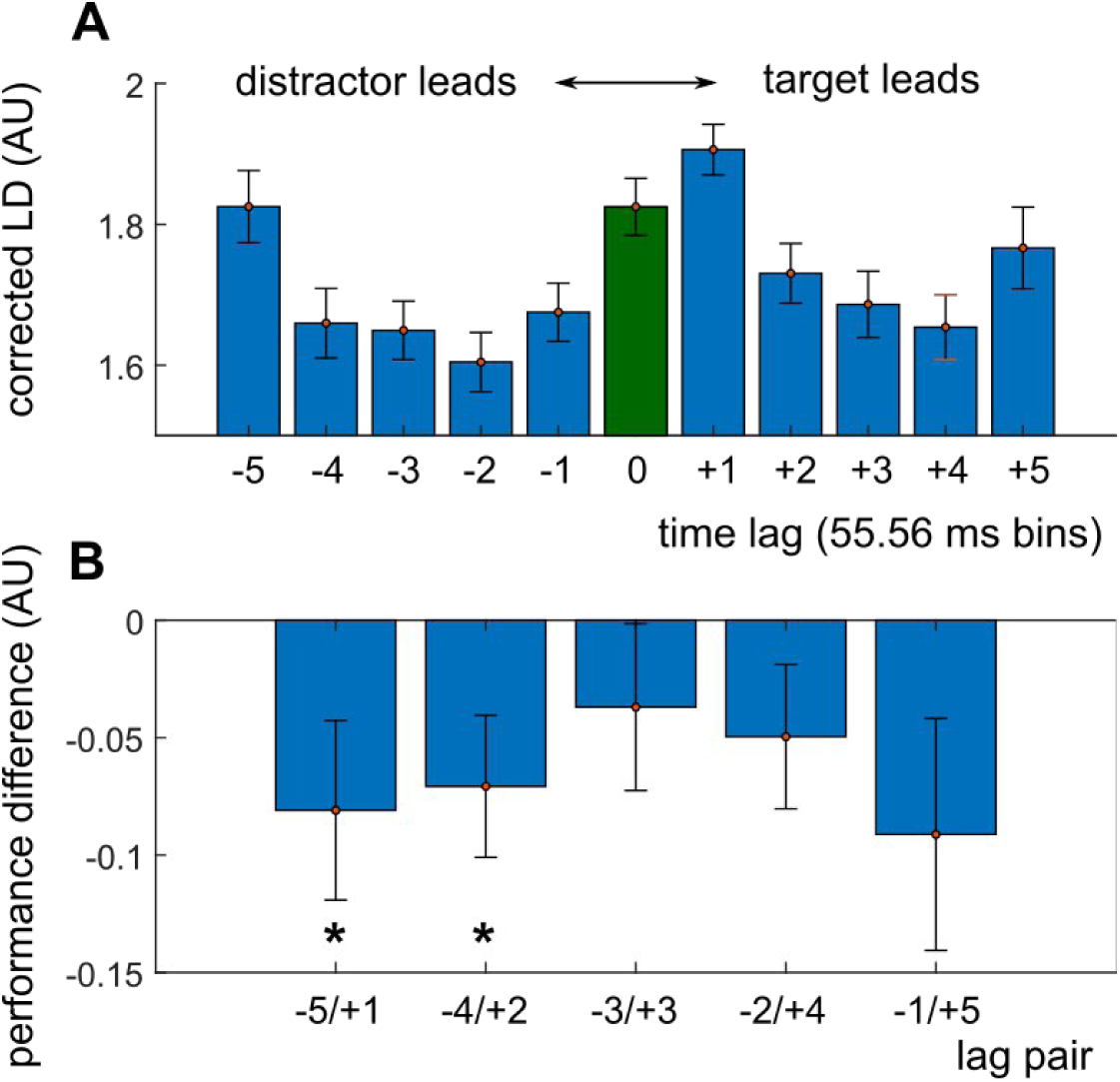
Acoustic/informational (A) and contextual (B) masking effects. **A**. Performance in the word report task as a function of time lag (expressed in bins of 55.56 ms) between the two speech streams. A negative lag indicates that the first word of distracting speech precedes target speech, the 0 (green bar) marks complete overlap. Due to the circularity in the design, the two streams are furthest apart at the −3/+3 lags. Performance is expressed as corrected Levensthein Distance (LD) (see Materials and Methods), i.e. higher values correspond to more errors. **B**. Performance difference for pairs of time lags where the middle three words in the two speech streams have the same lag, but that differ in which of the streams (target or distractor) starts earlier. Asterisks indicate a reliable difference from 0 (t-test). Error bars depict the standard error of the mean (SEM).

We hypothesized opposite best tACS phases when applied to modulate target or distracting speech, respectively. As an optimal (amplifying) phase for target speech should enhance speech perception, the same phase should disrupt speech perception for distracting speech, as distracting information is amplified (Keshavarzi et al., 2021). However, as the distribution of the difference between best tACS phases for target and distracting speech showed no significant deviance from uniformity (Figure 2B), testing for specific phase relationships (including the hypothesized phase opposition) was of no analytical use and not pursued further.

The methods described above quantify phasic modulation of speech perception, produced by tACS, but remain ignorant about the direction of the effect. It is thus unclear whether speech perception was improved, impaired, or both. As word report accuracy during tACS was expressed relative to that obtained during sham stimulation, positive and negative accuracy values reflect enhanced and impaired speech perception, respectively. However, as participants differed in their best (and worst) tACS phases, group-level statistics required the selection of individual tACS phase bins that led to most (and least) accurate perception, respectively. Such a selection leads statistical bias (Asamoah et al., 2019a); for instance, accuracy at the individual “best” phase is per definition a maximum, and likely to be positive (i.e. better than sham). We therefore repeated the selection procedure in the permutated datasets, described above, by extracting their best and worst tACS phases. This allowed us to quantify changes in accuracy that are due to the selection of behavioural extrema rather than reflecting a genuine tACS effect (grey line in Figure 3A). We defined enhancing tACS effects as the positive divergence from mean accuracy in the permuted dataset at the aligned best tACS phases, and disrupting tACS effects as the negative divergence at the aligned worst phase, respectively. We then contrasted the magnitude of enhancing and disruptive effects, and their interaction with condition (target vs distractor) and group (auditory vs IFG vs cutaneous) by means of a mixed (within-between) ANOVA.

**Figure 3.**
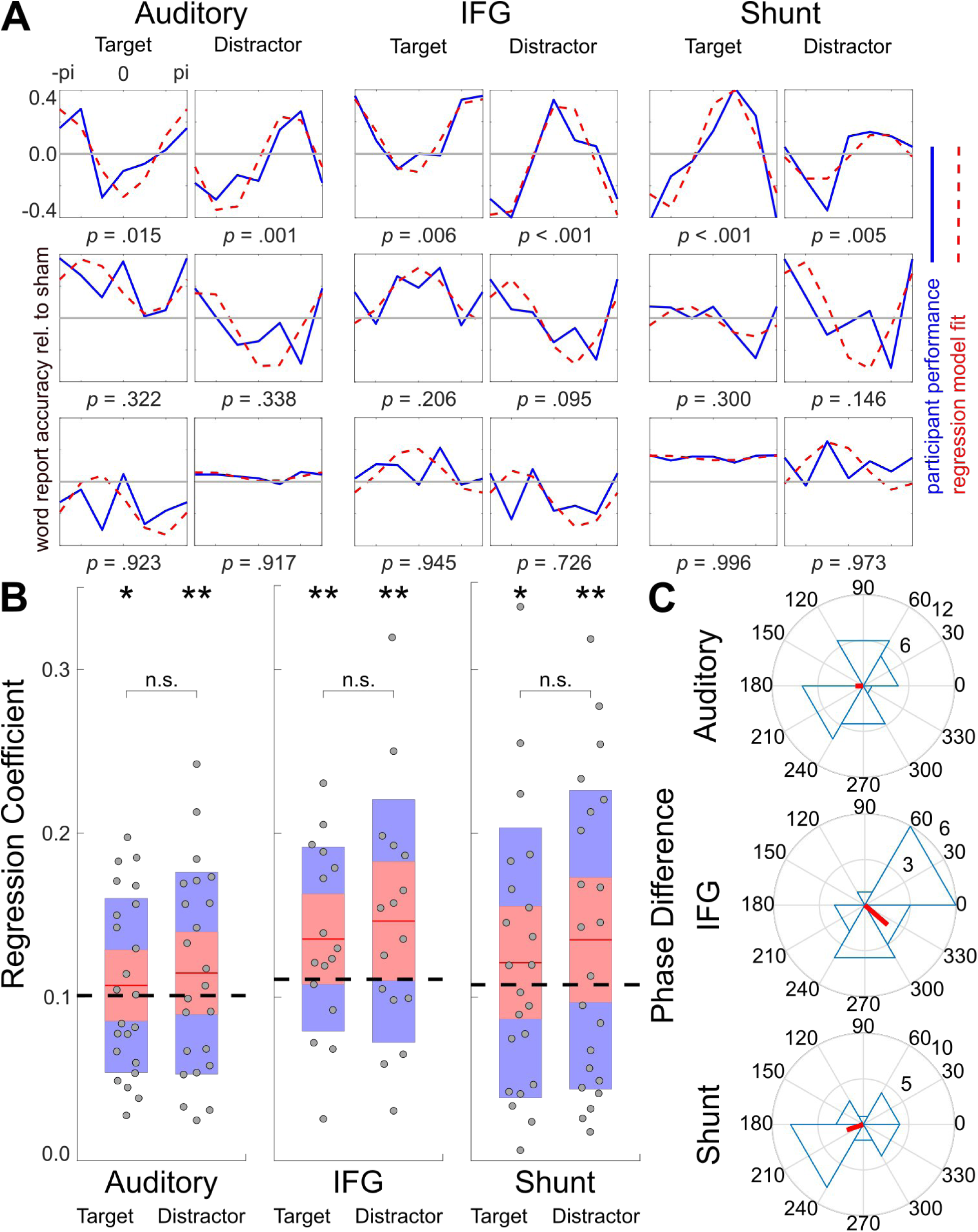
**A**: Word report accuracy as a function of tACS phase for nine example participants (blue line). For each montage and condition, participants with the best (top) median (middle) and worst (bottom) fitting regression models are shown. Accuracy is expressed relative to sham stimulation which included the same relative delays between target and distracting speech but no tACS. The red dotted line shows the fit of a sine function to the data. Its amplitude is equivalent to coefficients obtained from regressing accuracy on tACS phase (sine and cosine). These coefficients were contrasted with an intercept-only regression model to obtain individual p-values indicated below each panel. The peak of the sine fit was defined as individual participants’ best phase. Note that, for visualization of the phasic effect only, the +/-pi bin was plotted twice. **B**: Individual regression coefficients (root-mean square of sine and cosine coefficients) for target- and distracting speech and their distributions. Points, red lines, red areas and blue areas correspond to individual participants, their average, standard deviation, and 95% confidence interval, respectively. Dashed line indicates the 95% confidence interval of the means of the data from a surrogate distribution (generated separately for each dataset). Asterisks indicate average regression coefficients that exceed the upper limit of this confidence interval, reflecting a phasic modulation of word report accuracy. **C:** Distribution of circular differences between individual best tACS phases for the modulation of target speech and distracting speech. The direction and length of the red line illustrate the average difference and its consistency across participants, respectively.

## Results

### Effects and control of acoustic and contextual masking in speech perception

We asked participants to type in words from target speech, presented rhythmically at different phases of tACS (Figure 1A) or during sham stimulation. In three participant groups (one per stimulation montage), we used tACS to target auditory regions or IFG, or applied cutaneous stimulation (Figure 1C-D). Each participant received both active stimulation (auditory, IFG or cutaneous, depending on the group) and sham stimulation, albeit in different experimental blocks.

As expected and illustrated in Figure 2A, the time lag between target and distracting speech strongly influenced the accuracy of reporting target speech (F(10,50) = 10.55, p < .001, within-subject factor “time lag” in mixed ANOVA). However, as all possible lags between the two speech streams occurred equally often for all tACS phases, this effect of *acoustic/informational masking* (change in speech perception caused by the time lag between target and distracting speech) was counterbalanced across tACS phases in all stimulation and sham conditions. Indeed, acoustic/informational masking did not differ between stimulation and sham conditions (F(1,59) = 0.89, p = .35; within-subject factor in mixed ANOVA), nor between montage groups (F(2,58) = 2.06, p = .14; between-subject factor; no significant interactions between time lag, stimulation/sham, and montage). This result also suggests no generic effect of tACS (independent of phase) on speech perception.

Nevertheless, an inherent feature of our experimental design is that the target speech starts slightly earlier than the distracting speech in most trials for some tACS phases and slightly later for others. We illustrate this effect of *contextual masking* (changes in speech perception caused by the order of target and distracting speech) in Figure 2B, contrasting pairs of time lags where the middle three words in the two speech streams have the same lag but pair members differ in which of the streams (target or distractor) starts earlier (indicated by the sign in Figure 2). We found that contextual masking influenced the accuracy of reporting target speech, as some of the pairs reliably differed from 0 (p < 0.05 in a t-test against 0 for the pairs marked with an asterisk in Figure 2B). However, contextual masking – but not the effect of tACS – was also present during sham stimulation. Contextual masking did not differ between stimulation and sham conditions (F(1,59) = 0.40, p = .53; within-subject factor in mixed ANOVA), nor between montage groups (F(2,58) = 0.89, p = .42; between-subject factor; no significant interactions between pair, stimulation/sham, and montage). By expressing participants’ word report accuracy relative to that during sham stimulation, we were therefore able to control for contextual masking and isolate tACS phase effects (see Materials and Methods).

### tACS and cutaneous stimulation cause similar subjective sensations

To test whether skin sensations caused by electric stimulation was comparable between montage conditions, participants were asked to rate the intensity of their sensations at the end of each block (analysed on a scale between 0 = ”no sensations” and 10 = ”strong sensations”). Participants’ reported sensations did not differ between stimulation montages (average scores for Auditory montage: 2.22; IFG montage: 3.10; Shunt montage: 3.36; F(2,53) = 0.97, p = .39; between-subject factor in mixed ANOVA), nor between stimulation and sham conditions (F(1,54) = 1.3, p = .26; no significant interaction between montage and condition). In addition to near-identical electric fields in the scalp (Figure 1D), this finding speaks against one of the stimulation montages being less perceptible to participants.

### tACS and cutaneous stimulation cause similar phase effects in speech perception

We defined individual “best” tACS phase for speech perception as the peak of a sine function fitted to phase-resolved task performance (dotted red line in Figure 3A). In none of the three groups, the distribution of these best phases showed significant deviation from uniformity, neither for tACS applied to modulate target speech, nor for tACS applied to distracting speech (all p > 0.05; Rayleigh’s test). This result replicates that there was no consistent best phase across participants, as described previously (Neuling et al., 2012; Riecke et al., 2015; Zoefel et al., 2018; van Bree et al, 2021).

We then used linear regression with circular predictors to quantify whether tACS phase, relative to target or distracting speech, predicts word report accuracy of individual participants in the three experimental groups (see Materials and Methods). Figure 3B shows individual regression coefficients from this analysis (total of 61 participants across the three groups) and measures of their distribution. These coefficients are equivalent to the amplitude of the sine fits shown in Figure 3A and reflect how strongly speech perception was modulated by tACS phase. To assess statistical significance, we contrasted the obtained regression coefficients with those from another regression model that only uses the intercept (but no phase) to predict the same behavioural outcomes (F-test), yielding one p-value per participant. Individual p-values were combined to a single group-level p-value per condition with Fisher’s method. This group-level p-value (rather than individual p-values) was used to draw statistical conclusions. This approach is sensitive to phase effects and does not produce more false positive results than expected from the significance threshold used (5%) (Zoefel, Davis, et al., 2019).

We found that tACS over auditory regions reliably modulated word report accuracy, irrespective of whether it was applied to alter neural tracking of target speech (*p* = .016) or neural tracking of distracting speech (*p* = .0008). We observed a similar phasic modulation for tACS of IFG (target: *p* = .0006, distractor: *p* < .0001) and importantly, for cutaneous stimulation (target: *p* = .0002; distractor: *p* < .0001). The dashed black lines in Figure 3B illustrate the upper limit of a 95% confidence interval for regression coefficients obtained under the null hypothesis (from permuted datasets; see Material and Methods). The fact that the mean regression coefficients in all conditions (red lines in Figure 3B) exceed the confidence interval confirms the presence of tACS phase effects in all participant groups.

When tACS phase relations to both target and distracting speech were used in combination to predict behavioural outcomes, the accuracy of the regression model increased further, and this improvement was reliable across stimulation montages (Mixed ANOVA using Fisher’s z to contrast regression coefficients from single vs combined models, main effect of model F(2,58) = 10.51, *p* = .0001, no effect of montage: F(2,58) = .85, p = .434, no interaction: F(4,56) = .38, p = .834). This result indicates that neural tracking of both target and distracting speech jointly influenced speech perception.

We found no difference in tACS phase effects on speech perception between target/distractor conditions (*F*(1,59) = .18, *p* = .677; within-subject factor in mixed ANOVA on regression coefficients) or montages (*F*(1,59) = .23, *p* = .636; between subjects), and there was no interaction (*F*(3,57) = .8, *p* = .500). During cutaneous stimulation, the electric field in the brain was reduced but not zero (Figure 1D), and it is possible that the residual cortical stimulation (rather than cutaneous stimulation) produced the phasic modulation of speech perception. However, in such a case the effect would be expected to be reduced as compared to a montage with equivalent cutaneous stimulation, but stronger cortical stimulation. We therefore compared the magnitude of the phasic effect (i.e. the regression coefficients shown in Figure 3B) in the cutaneous stimulation group with that in the auditory group, reduced in proportion (shunt = ∼50% reduced cortical stimulation) to the change in cortical electric field. We found that the phasic modulation during cutaneous stimulation was significantly stronger than that expected from remaining cortical stimulation, given the reduction in cortical electric field (attended: t(43) = −5.56, p < .0001, t(43) = −4.97, distractor: p < .0001).

As explained above, the sham condition in our design does not act as a representation of the null hypothesis, but controls for contextual masking. Instead, the null hypothesis is represented by the intercept-only model used for statistical analysis, and by the confidence intervals for regression coefficients obtained from permuted datasets. Nevertheless, the fact that the phase effect in all stimulation conditions is statistically reliable after subtraction of the sham data can only be explained by a stronger phase effect in these conditions than during sham stimulation.

Together, we found a phasic modulation of word report accuracy in all groups, and no difference between them. As groups differed in their amount of brain stimulation but received similar cutaneous stimulation (Figure 1D), this pattern of results shows that the cutaneous stimulation that goes along with conventional tACS may cause phase effects in speech perception. Nevertheless, the phasic effects also suggest a causal role of neural tracking for speech perception in multi-speaker settings.

### Best tACS phases for target and distractor modulation are uncorrelated

The optimal (amplifying) tACS phase for target speech should disrupt speech perception when applied to distracting speech as distracting information is amplified, predicting opposite best tACS phases for target and distracting speech, respectively (Figure 1B). However, the distribution of the difference between best tACS phases for target and distracting speech showed no significant deviance from uniformity (Figure 3C; Auditory: p = .681, z = .391, IFG: p = .176, z = 1.75, Shunt: p = .283, z = 1.27). Given the uniform distribution, testing for specific phase relationships (including the hypothesized phase opposition) was not meaningful and not pursued further. This finding contradicts our hypothesis that best tACS phases for target and distractor modulation are in anti-phase.

### tACS enhances and disrupts speech perception

We examined whether the phasic effects we observed corresponded to enhanced or disrupted speech perception (or both), respectively. Figure 4A shows average word report accuracy (relative to sham) after re-alignment to individual best tACS phases in each condition and group (see Materials and Methods). This analysis was not designed to assess phasic effects (this was already done in analyses described above), but to contrast enhancing- and disrupting effects on perception. Due to the alignment, the behavioural data necessarily follows a sinusoidal shape, and behavioural extrema are located at the aligned best phase and its anti-phase, respectively. We illustrate how much of this is due to the re-alignment itself by showing the outcome from the same procedure in a permuted dataset (grey line in Figure 4A). Note that the actual data diverges from the simulated null effect in both directions (positive and negative). We first verified that the sinusoidal shape in the permuted datasets is due to the re-alignment itself rather than to the presence of phasic effect in the permuted datasets. For each montage and condition, we counted the proportion of significant phase effects (group level p < 0.05) in the 1000 permutations. We found that this proportion to be close to 5% (range between 4% and 5.5 %), as expected from the significance threshold of α = 0.05. This confirms our previous result that the regression- and permutation-based analysis does not produce more false positive results than expected (Zoefel et al., 2019). We then defined enhancing tACS effects as the positive divergence from the null effect at the aligned best tACS phases, and disruptive tACS effects as the negative divergence at the aligned worst phase, respectively.

**Figure 4.**
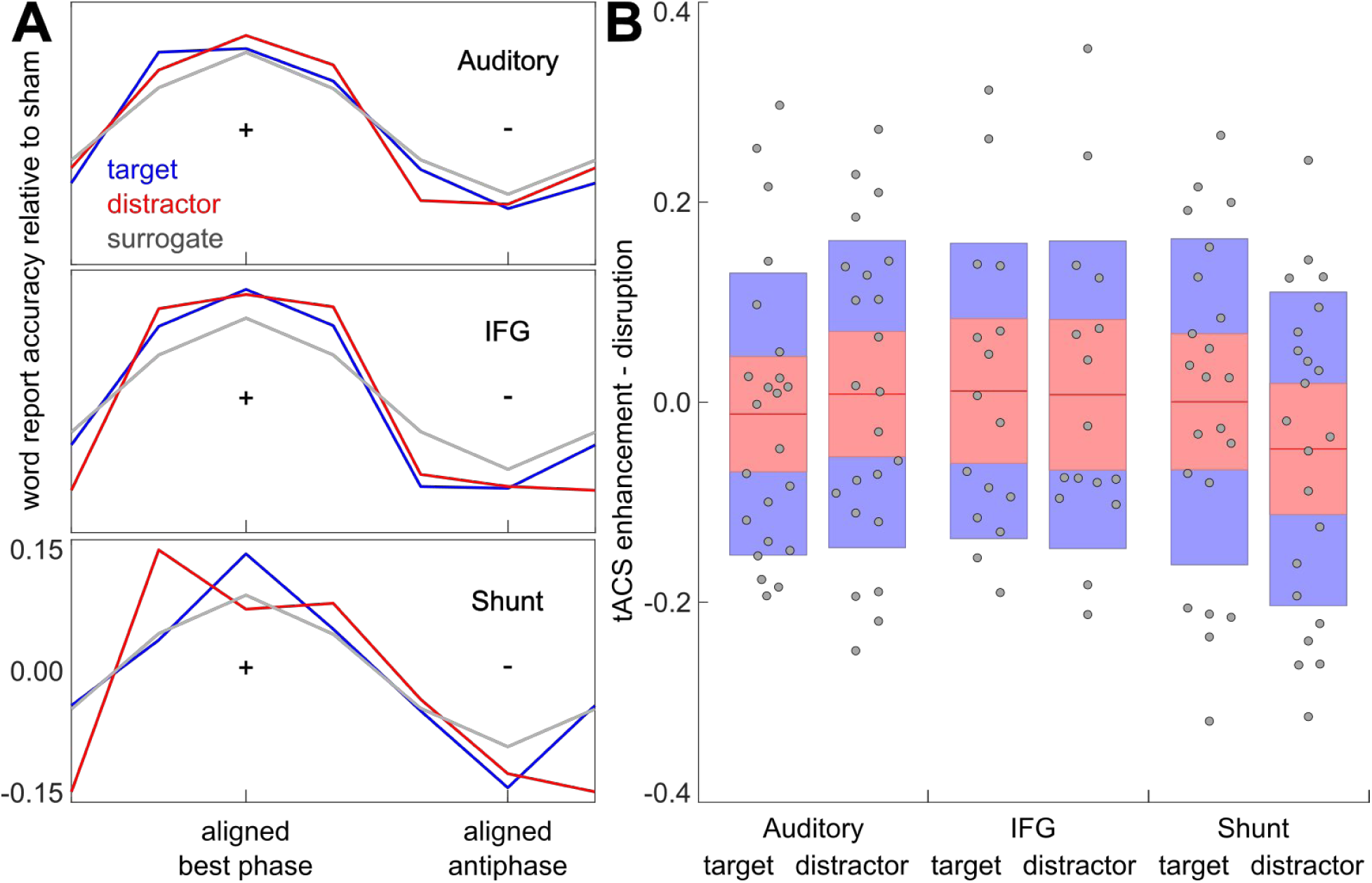
**A**: Averaged word report accuracy (relative to sham), re-aligned so that individual best tACS phases (peak of fitted sine function) fall into a common phase bin. Re-alignment was done separately for each of the three montages and the two tACS conditions, as well as their respective null/surrogate distributions. For the latter, the average across 1000 permutations is shown. **B**: Enhancing and disrupting effects of tACS, expressed as changes in accuracy (relative to surrogate average) at the best (+ in A; peak of fitted sine) or worst (- in A; trough of fitted sine; multiplied by −1 for comparability) tACS phase for word report accuracy, respectively. Points, red lines, red areas and blue areas correspond to individual participants, their average, standard deviation, and 95% confidence interval, respectively.

Differences between enhancing and disruptive tACS effects are shown in Figure 4B. For none of the target/distractor conditions or stimulation montages, a significant difference between enhancement and disruption was apparent (main effect enhancement vs disruption: *F*(1,59) = .14, *p* = .710; no significant interaction with condition or montage in mixed ANOVA). Given the significant phasic effects reported above, this result indicates that tACS led to *both* an enhancement and disruption of speech perception, depending on its phase relation to the speech stimulus.

## Discussion

In three experimental groups, we used tACS to independently manipulate neural activity aligned to two simultaneously presented speech streams. We find that (1) neural tracking of both target- and distracting speech causally contributes to speech perception; (2) these effects are of similar magnitude and (3) their combined effect goes beyond each of the two individual modulations alone. However, when comparing results from three different stimulation montages, including a control condition with similar cutaneous stimulation but significantly less brain stimulation, we found comparable effects. This implies that, although tACS effectively modulated speech perception, cutaneous stimulation produced by tACS can cause similar phase effects.

### Cutaneous tACS effects on speech perception need to be controlled for

Most of the current applied during tACS is shunted by the skin (Antal & Herrmann, 2016; Neuling, Wagner, et al., 2012), leading to relatively strong cutaneous- and relatively weak brain stimulation, respectively. A significant challenge for tACS research is therefore a potential stimulation of peripheral nerves and its effects on the brain. Indeed, Asamoah et al. (2019b) showed that the application of topical scalp anaesthesia under tACS electrodes significantly reduced the entrainment of neural activity in the motor cortex to tACS.

Although a role of cutaneous stimulation for tACS effects in other modalities than the motor system seems possible, it is rarely tested or controlled for. This includes the field of speech perception where tACS has become a popular tool (Erkens et al., 2020, 2021; Kadir et al., 2020; Keshavarzi et al., 2020, 2021; Kösem et al., 2020; Marchesotti et al., 2020; Meier et al., 2019; Preisig et al., 2020, 2021; Riecke et al., 2015; Rufener, Oechslin, et al., 2016; Rufener et al., 2019; Wilsch et al., 2018; Zoefel et al., 2020), due to the prominent role of neural tracking in speech processing (Di Liberto et al., 2015; Ding et al., 2014; Ding & Simon, 2014; Myers et al., 2019) and tACS’ ability to manipulate it (Riecke et al., 2018; Wilsch et al., 2018; Zoefel et al., 2020). As tACS-induced sensations are maximal at the onset and offset of the stimulation (Thiele et al., 2024), a typical sham condition mimics these sensations and consists of only brief stimulation at the beginning and end of each experimental block (Schutter & Wischnewski, 2016; Thiele et al., 2024). Other studies used a different tACS frequency as a control measurement (Rufener, Oechslin, et al., 2016; Rufener, Zaehle, et al., 2016), but these studies were based on a modulation of power (rather than phase) of oscillatory activity, and sensations (and possibly cutaneous stimulation) seem to differ between tACS frequencies (Turi et al., 2012, 2013). Consequently, none of these controls can provide an unambiguous differentiation between effects of cutaneous stimulation and those of direct cortical stimulation. Using an recently proposed approach to control for cutaneous stimulation (Neri et al., 2020), we here show that such stimulation can indeed cause phasic effects on speech perception resembling those produced by more conventional tACS montages. Such a modulation might stem from rhythmic activation of somatosensory pathways which is then, via the well-described connectivity between somatosensory- and auditory systems (Caetano & Jousmäki, 2006; Gick et al., 2010; Kayser et al., 2005; Lakatos et al., 2007; Riecke et al., 2019), relayed to the auditory cortex to entrain neural activity there. This notion is supported by a previous demonstration that the speech envelope can be conveyed via vibrotactile stimulation to modulate speech perception (Guilleminot et al., 2023; Guilleminot & Reichenbach, 2022). We hypothesize that the cutaneous stimulation produced by tACS acts in a similar way.

Additional evidence for a separate cutaneous pathway during tACS is the similarity in effect size for target- and distractor manipulations in the present study, even when targeting IFG. As IFG is higher in the auditory hierarchy where neural representations of targets and distractors are more distinct (Zion Golumbic et al., 2013), we expected to find similarly distinct effects of tACS on targets and distractors, but this hypothesis was not confirmed. Our data might involve a general effect of cutaneous stimulation that is independent of the site of stimulation, and therefore modulates perception irrespective of the target.

Our results do not suggest that all effects of tACS can be explained by cutaneous stimulation. Single-neurons recordings in non-human primates show that tACS still entrains neural activity when somatosensory input is blocked (Vieira et al., 2020). Moreover, tACS aftereffects correlate with the cortical electric field rather than with the scalp field (Kasten et al., 2019), and the magnitude of tACS effects in magnetoencephalography (MEG) and functional brain imaging (fMRI) correlates with the electric field strength produced (Kasten et al., 2019; Preisig & Hervais-Adelman, 2022). Our results thus do not discredit the notion of cortical entrainment through tACS, but rather illustrate the urgent need for control montages to separate direct cortical-from cutaneous effects, especially when targeting specific regions.

### Neural tracking at the “cocktail party”

Irrespective of the underlying mechanism – we here show that a manipulation of neural tracking changes perception of speech at the “cocktail party”. Recent research has examined neural dynamics (including neural tracking and entrainment) in such multi-speaker scenarios. Orf et al. (2022) compared EEG “tracking” of target- and distracting speech with that of a neutral stimulus. Although all speech stimuli evoked tracking responses, only the target speech produced clear differences to the neutral speech. This implies that neural tracking of attended speech is functionally more relevant than that of distractors, contradicting our result that the causal contributions of target- and distractor tracking to speech perception are of similar magnitude. However, it is possible that the tracking of distractors is particularly relevant in challenging situations: Fiedler et al. (2019) found that distracting speech evokes a late tracking component that is of opposite polarity to that seen for target speech, but only when it is difficult to understand, implying an additional, active suppression of distractors. In line with this notion, the task used by Orf et al (2022) involved spatial separation cues, which are known to be important for speech segregation (Shinn-Cunningham, 2005; Shinn-Cunningham & Best, 2008). As in our study, both speakers were presented binaurally through in-ear headphones, cortical tracking of distracting stimuli might have become more relevant due to the increased difficulty in absence of these cues.

Our study is closely related to that by Keshavarzi et al. (2021), who applied tACS to independently modulate tracking of target- and distracting speech, and found effects on speech perception in both cases. Although our results confirm these findings, we did not replicate the anti-phase relationship between target- and distractor tracking that was apparent in their work. In the study by Keshavarzi and colleagues (2021), delaying the theta-filtered envelope with tACS might not only have altered theta-neural activity relative to the acoustic stimulus, but also de-synchronized theta activity with that at slower time scales (e.g., delta), which also plays a role for speech perception (Ding & Simon, 2014; Etard & Reichenbach, 2019; Keitel et al., 2018; Meyer et al., 2016; Molinaro & Lizarazu, 2018). Stimulation in the delta band may be a promising control in future work, given that its effect on speech perception can be dissociated from those of theta-tACS (Keshavarzi et al., 2020). In our study, we used rhythmically spoken and unrelated mono-syllabic words instead of natural speech. This manipulation avoided the concomitant fluctuations in information at multiple times scales that natural speech has (Doelling et al., 2014; Ghitza, 2012; Giraud & Poeppel, 2012). Moreover, whilst our tACS protocol manipulated tracking at the word/syllable level, stimuli used by Keshavarzi et al. (2021) included grammatical- and structural information. We speculate that the paradigm used by Keshavarzi et al. (2021) induced competition between target and distracting speech (and opposite best tACS phases) by manipulating or de-synchronising entrained activity at multiple hierarchical levels. In our study, best tACS phases for target- and distracting speech were uncorrelated, which implies that their processing involves distinct neural mechanisms, at least at lower hierarchical levels.

### Implications for clinical applications

It has been proposed that tACS can have clinical applications, specifically when applied to improve speech-in-noise comprehension (e.g., Erkens et al., 2021; Keshavarzi & Reichenbach, 2020; Riecke et al., 2015). Ageing and hearing-impaired listeners do not only struggle to understand speech in noise, but also show changes in neural speech tracking, implying a functional role for speech perception in noisy scenarios (Cabeza et al., 2002; Henry et al., 2017; Petersen et al., 2017). It is a common complaint amongst users that hearing aid can be inefficient in complex hearing scenarios (Fischer et al., 2020; Kochkin, 2000), and speech stream segregation worsens with age (Getzmann et al., 2017; Getzmann & Näätänen, 2015; Pichora-Fuller et al., 1995; Pichora-Fuller & Singh, 2006). Our findings can also be seen in such a context, where tACS can possibly assist struggling listeners by manipulating neural tracking in such situations. However, apart from obvious methodological and technical challenges that remain to be solved, these findings also imply that tACS’ clinical usefulness to improve speech perception should be compared with alternative methods: For example, a rhythmic tactile stimulus might achieve the same purpose with much simpler means (Guilleminot et al., 2023; Guilleminot & Reichenbach, 2022).

We found that the simultaneous manipulation of neural tracking of targets and distractors goes beyond what can be achieved with the manipulation of either of them alone. This finding is important as reported tACS effects are typically small (Bland & Sale, 2019; Riecke, 2016; Riecke & Zoefel, 2018) – indeed, effects of acoustic or informational masking (Figure 2A) on speech perception seemed stronger than those of tACS in our study (although a systematic comparison between the two require a different experimental approach). Approaches to enhance tACS effects to clinically relevant levels are therefore needed. Our design required rhythmic speech to examine these combined effects in the same trials. An important step would therefore be the development of a stimulation protocol to simultaneously modulate target- and distracting speech in natural scenarios.

## Conclusion

We here demonstrate that neural tracking of target- and distracting speech jointly modulates speech perception in a multi-speaker scenario. Nevertheless, a montage with similar cutaneous stimulation but reduced brain stimulation caused similar phasic effects in word report accuracy, possibly by entraining auditory brain regions through connectivity with somatosensory regions. These results illustrate the urgent need for montages controlling for cutaneous stimulation in future tACS work on speech perception and beyond.

## Data and Code Availability Statement

Data and code will be made available upon request to the authors.

## Acknowledgements

This work was supported by the Fondation pour l’Audition (grant number FPA-RD-2021-10) and by EUR CARe (N°ANR-18-EURE-0003) in the framework of the Programme des Investissements d’Avenir.

